# Foraging competence and scrounging tolerance enhance social relationships in a socially tolerant wild primate

**DOI:** 10.1101/2025.06.12.659242

**Authors:** Elif Karakoc, Richard Vogg, Michele Marziliano, Jacob Petersdorff-Campen, Alexander Ecker, Peter M. Kappeler, Claudia Fichtel

## Abstract

Social interactions are crucial for individual health and ultimately fitness, making the choice of social partners particularly important. Previous research has shown that individuals who succeed in foraging tasks often receive increased affiliation from group members. Similarly, in a social learning context, individuals who possess valuable information become more attractive social partners. Thus, an individual’s role in a foraging context–specifically, whether it is a successful producer–can influence its social relationships. Therefore, we examined the interplay between social learning, producing and scrounging behavior, and social relationships in four groups of wild redfronted lemurs (*Eulemur rufifrons*). We conducted an open diffusion experiment with food boxes that required animals to learn one of two techniques to open them. 27 out of 29 individuals participated in the experiment, 24 interacted with the boxes and 16 learned to open them. Initial success was better predicted by use of individual than social information, i.e., manipulating the food boxes vs. time spent watching successful individuals or scrounging. Older males were less successful than females. Scrounging occurred in about 26% of events, with on average 1.3 individuals scrounging. The technique used, age and sex of the producer did not predict scrounging frequency. Learners and males scrounged more often than non-learners and females. Among learners, less successful individuals scrounged more often and this effect was more pronounced in males. More successful individuals and those that were scrounged more often received more affiliative behavior. Thus, cognitive skills and scrounging tolerance may strengthen social relationships in this primate species.

## Introduction

Affiliative social interactions can have far-reaching effects on an individual’s health and ultimately fitness, such as higher reproductive success, greater offspring survival and improved longevity (Snyder-Mackler et al. 2020). Natural selection therefore favors individuals that have the ability to effectively compete and cooperate with others, which in turn requires the ability to assess other’s motivations, intentions and skills (Seyfarth and Cheney 2015; Cheney et al. 2016). In species that live in large, permanent and cohesive groups, where known individuals interact repeatedly with each other, individuals do not interact randomly with each other (Seyfarth and Cheney 2012; Ostner and Schülke 2018). As a result, emergent dyadic social relationships vary qualitatively in several dimensions (Silk et al. 2013), creating differentiated social networks (Lusseau and Newman 2004; Farine and Whitehead 2015). While earlier research identified factors such as kinship, cooperation or phenotypic assortment predisposing some individuals to be more valuable social partners than others (Seyfarth and Cheney 2012), more recent work has suggested that specific skills may make some individuals preferred social partners (Fruteau et al. 2009; Kulahci et al. 2018; Kulahci and Quinn 2019).

Experimental studies with long-tailed macaques *(Macaca fascicularis),* vervet monkeys (*Chlorocebus pygerythrus)*, and Guinea baboons (*Papio papio)* revealed that low-ranking individuals knowing how to manipulate a food dispenser, providing food for the entire group, received in turn more affiliative interactions from other group members (Stammbach 1988; Fruteau et al. 2009; O’Hearn et al. 2025). Hence, these non-human primates are able to monitor and recognize the expertise of others and adapt their own behavior in a fine-tuned manner to take advantage of these capabilities (Stammbach 1988). In a similar vein, a study on ring-tailed lemurs *(Lemur catta)* showed that individuals in the vanguard of adopting an innovation that allowed them to open a food dispenser also received more affiliation, even though – crucially – no other group member enjoyed a personal food reward (Kulahci et al. 2018). This study suggested that the mere possession of information or being competent can increase an individual’s value as a social partner.

Competence refers to the skills required in a particular domain, such as foraging, and reflects an individual’s overall ability to succeed in a given domain (Sih et al. 2019). In the context of social learning, competent individuals, i.e. innovators of a new behavior and thus successful individuals, play a key role in information transmission, as interacting with successful individuals can lead to the learning of new skills by others. Social learning is defined as changes in an individual’s behavior that result from exposure to another individual’s behavior or its products (Heyes 1994). Social relationships, in turn, are crucial for the transmission of information within a group and act as channels for information exchange (Hoppiq et al. 2010, 2011).

Studying innovation and information transmission in natural serngs is diffcult because the likelihood of witnessing and documenting this process is rather low (Whiten and Mesoudi 2008; Schnoell and Fichtel 2012; Fichtel 2022). To overcome this challenge, many researchers have conducted social diffusion experiments (Whiten and Mesoudi 2008; Whiten et al. 2016). These experiments typically present a food box that can be opened by using one or more techniques. In one variant, the social diffusion paradigm, a demonstrator can be trained in advance to subsequently document the diffusion of the opening technique within a group. In another variant, the open diffusion paradigm, no demonstrator is trained in advance and individuals can simultaneously explore and learn individually how to solve the task (Canteloup et al. 2020, 2021; Barreq et al. 2017; Arbon et al. 2023). In particular, open diffusion experiments enable the investigation of characteristics of innovators who acquire primary knowledge in a natural way that is not influenced by training (Whiten and Mesoudi, 2008). Another advantage of open diffusion experiments is that naïve individuals are not seeded with a trained technique. Therefore, the natural preference and flexibility towards the solutions can be observed (Arbon et al. 2023).

In social learning experiments in which several individuals can interact at the experimental apparatus, it has been documented that naïve individuals exhibit scrounging behavior, i.e., benefirng from the efforts of others rather than actively learning by themselves (e.g. Fragaszy and Visalberghi 1989; Beauchamp and Kacelnik 1991). The effect of scrounging in the context of social learning is controversial. Some studies suggest that scrounging tended to inhibit social learning (Giraldeau and Lefebvre 1987; Beauchamp and Kacelnik 1991) or learning in general (Aplin and Morand-Ferron 2017; Reichert et al. 2021; Lefebvre and Helder 1997). These studies suggested that scroungers who rely on the efforts of others rather than learning for themselves may be hindering their own acquisition of skills and competence. However, Caldwell and Whiten (2003) showed that scrounging from competent individuals can facilitate learning compared to simply observing competent individuals. This suggests a complex interaction in which scrounging may serve not only as a passive behavior, but also as an active mechanism that positively contributes to the learning experience within social groups. Therefore, the role of scrounging in social learning contexts needs to be investigated more comprehensively, in particular in natural serngs.

Observations that competent individuals become more valuable social partners when they possess new information (Kulahci et al. 2018) or provide food that enables others to learn socially or scrounge (Stammbach 1988; Fruteau et al. 2009; O’Hearn et al. 2025), raise the question of whether affiliation could be traded as a commodity for gathering new information or scrounging. Therefore, we investigated in this study the interplay between social learning, scrounging, and social relationships in wild redfronted lemurs (*Eulemur rufifrons*). Understanding how individuals navigate complex social interactions and leverage social information can provide insights into the evolutionary significance of behavioral flexibility and social competence (Kulahci and Quinn 2019). Lemurs (Lemuriformes), an adaptive radiation of primates endemic to Madagascar, are interesting in this context because they evolved in isolation from other primates for more than 50 Myr (Yoder et al. 1996) and group-living has evolved twice independently (Kappeler and Pozzi 2019). They are also considered a model for early primate cognitive evolution (Fichtel and Kappeler 2012; Ho et al. 2021).

Redfronted lemurs, in particular, are an interesting study species to investigate the interplay between social learning, scrounging, and social relationships for several reasons. Firstly, redfronted lemurs have a relatively egalitarian social structure that promotes a high degree of social tolerance (Pereira et al. 1990; Ostner and Kappeler 1999; Fichtel et al. 2018), facilitating social learning (Coussi-Korbel and Fragaszy 1995; van Schaik et al. 1999). Secondly, in previous studies we have already investigated the cognitive abilities of redfronted lemurs in the wild, including social learning and innovation (Schnoell and Fichtel 2012; Schnoell et al. 2014; Huebner and Fichtel 2015; Fichtel 2022), and have documented scrounging behavior in social learning experiments using a social diffusion paradigm (Schnoell and Fichtel 2012). However, it remains unclear whether scrounging facilitates or inhibits social learning. Thirdly, redfronted lemurs are known to exchange grooming as a commodity for tolerance (Port et al. 2009), prompting the question of whether they also trade affiliative behavior for opportunities to scrounge or gain access to social information from skilled individuals.

To address these open questions, we conducted an open social diffusion experiment by presenting four groups of redfronted lemurs with food boxes that could be opened by using two different techniques (lifting or pushing) to gain access to a food reward (Figure 1a, b). Specifically, we studied the following questions: (I) Who innovates and do redfronted lemurs develop a preference for one technique over the other? (II) What predicts initial learning: Use of individual information by manipulating the food box or social information, i.e., watching successful individuals or scrounging? (III) Who scrounges from whom? (IV) Do competent individuals who are more successful at opening the food boxes or those who allow others to scrounge more often receive more affiliative behavior, thus becoming more valuable social partners?

**Figure 1:**
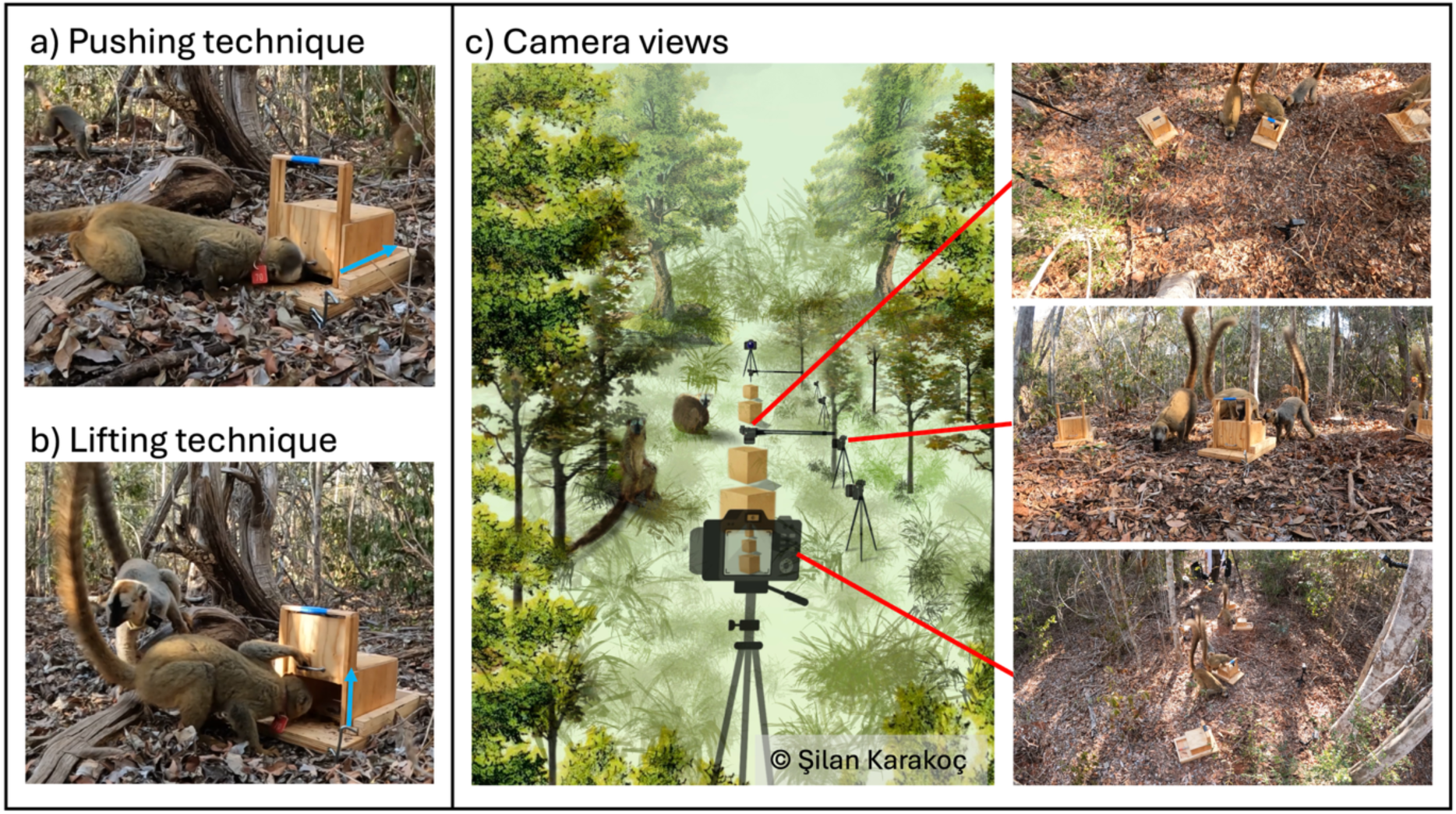
A redfronted lemur using a) the pushing and b) the lifting technique to open the food box. c) Demonstration of the experimental setup; an overview of the setup (center) and the camera views; top view can be seen on the upper right, close-up view is middle right, and the wide-angle camera view is bottom right.

**Figure 2:**
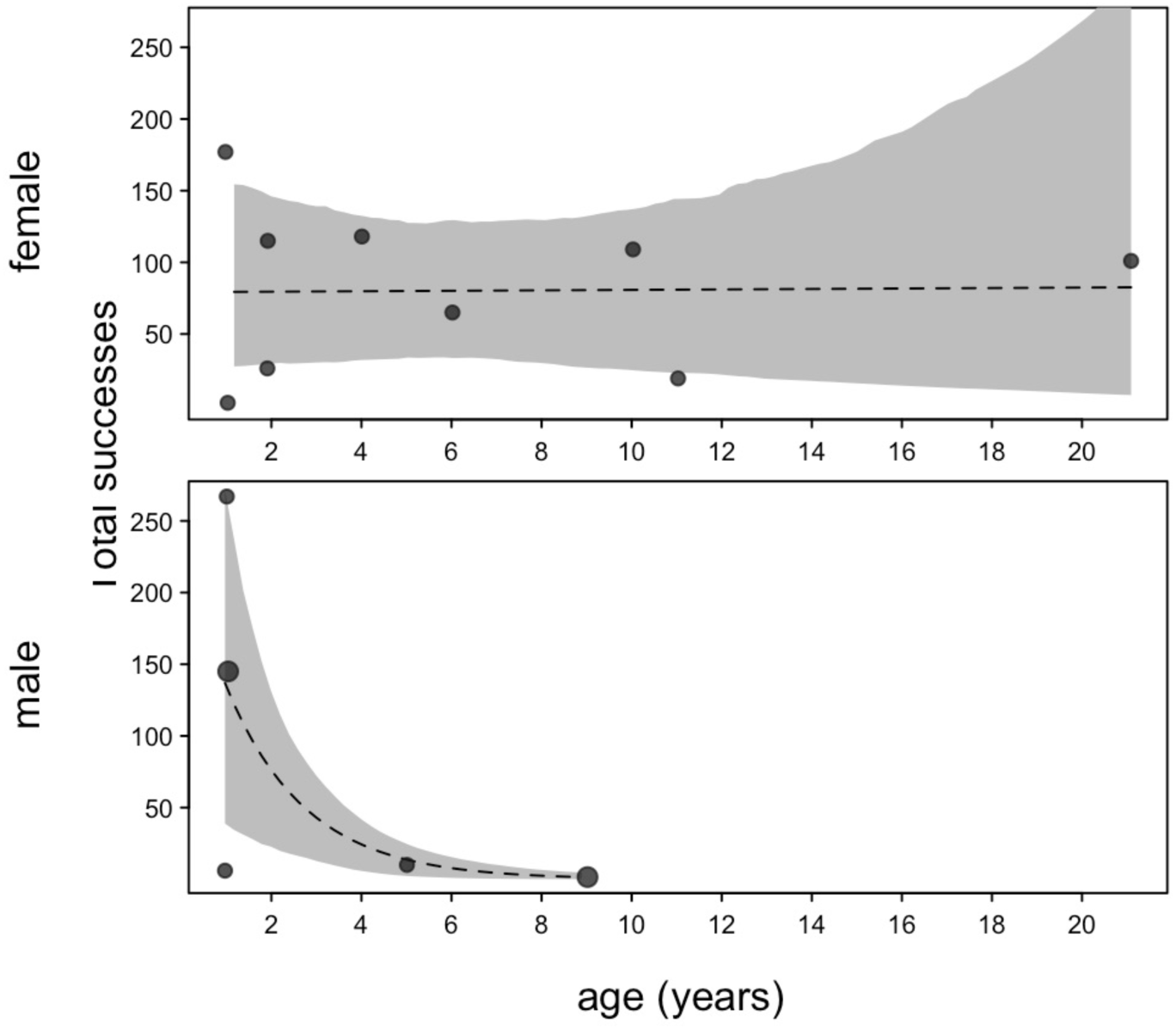
Depicts the effects of sex and age (in years) on the number of successful openings. Dashed lines indicate the regression lines and shaded areas the 95% confidence intervals.

## METHODS

### Field Station and Subjects

We conducted this study at the German Primate Center (DPZ) research station in Kirindy Forest, western Madagascar (Kappeler and Fichtel 2012). We conducted the experiments on four groups of red-fronted lemurs (*Eulemur rufifrons*), comprising a total of 31 individuals. One adult female from group B and one adult male from group R1 left their groups during the study period. As part of a long-term study (Kappeler and Fichtel 2012), the study animals are marked with a collar and a tag for individual identification and are well habituated to humans.

### Experimental procedure and setup

We used two-option-food-dispenser (20x20x30 cm) for this experiment (Figure 1a, b). The food box consisted of two parts: a wooden foundation with a cavity containing the food reward and an upper box composed of four square boards and a sliding door aligned in a cube-shaped manner. Individuals could access the food reward by lifting the sliding front-door or pushing back the box on the foundation (Figure 1). The sliding door would fall back shut if not held open. Similarly, if the box was opened by pushing, metal springs would return it to its initial position after release. Since both techniques were implemented to the same part of the food box, that is the door, simple social learning mechanisms like local or stimulus enhancement should not account for copying lifting or pushing (Huber et al. 2009; Schnoell and Fichtel 2012). In addition, both techniques were likely to have the same level of difficulty. The food box was filled with a mixture of raisins and pieces of peeled oranges as food rewards. At the beginning of each experiment, we set up the food boxes and the cameras at a distance of about 50 m from a study group to avoid animals witnessing experimenters filling the food boxes with the food rewards. Afterwards we attracted redfronted lemurs by presenting an acoustic signal produced by a clicker that is commonly used to train animals. To prevent animals from associating the presence of humans with food, experimenters additionally wore a white lab coat. Animals were first habituated to the new food boxes by presenting them with open or half-open food boxes containing the reward for three to four sessions. After the habituation phase, food boxes were presented with closed doors for 10 experimental sessions, depending on the group over a period of four to six weeks. Experimental sessions lasted between 10 and 30 minutes (mean ± SD: 13.8 ± 5.1 min). Sessions ended when all individuals lost interest and left the experimental arena for at least one minute.

### Video-recordings and temporal video synchronization

For each session, we set up four food boxes anchored to the ground and recorded the setup using eight GoPro Hero 10 cameras. These included four front-facing ground cameras, two overview cameras on tripods, and two tree-mounted cameras capturing top views (Figure 1 c). Although the cameras can be voice-controlled, the starting times were not completely synchronized. In some cases, we had to start them manually, when the voice recognition did not work for single cameras. As a result of the initial misalignment, the videos were temporally unsynchronized and had to be later synchronized. To this end, we started one of the cameras manually and turned off the voice control for this camera. After storing information about the date and the session, we started the other cameras using voice control and checked whether all cameras started recording; otherwise, we turned them on manually.

We used the clicker with an irregular audio pattern, both to attract the lemurs and to create a distinctive audio signal that we used for temporal alignment of the video-recordings. Using Python and the libraries wave and numpy (Harris et al 2020), we read in the audio files for each camera. Since the GoPros automatically cut a recording into chunks of 4GB, we stitched recordings together based on their file names to have one large recording for each camera and experiment. Afterwards, we converted the audio signal into a mono format to have one single channel of audio data, and for performance reasons sampled every 100th time step. Starting from the beginning of the main camera, which always started recording before all other cameras, we used a sliding window approach to find the offset for each other camera with respect to the main camera. For each potential offset, we calculate the absolute difference between the audio signal of the main camera with the audio signal of the other camera. This results in a time series of potential offsets and corresponding differences. The most distinctive local minimum of this time series corresponds to the best match of the audio signals of both cameras. We used the Python function find_peaks from the library scipy.signal (Virtanen et al. 2020) to detect this minimum. Finally, we added empty frames to the beginning of each video. The number of frames was the determined offset, and no frames were added to the main camera. This resulted in temporally synchronized videos, all videos had a resolution of 1920x1080.

### Video annotations

Synchronized video recordings were annotated using BORIS v.7.13.9 (Friard and Gamba 2016), scoring the following behaviors:

1. Manipulating the box (duration): touching the food box with either the hand or any part of the head for at least 1 s
2. Success (count): successful opening of the food box with the individual gaining access to the food reward
3. The technique used to open the food box
4. Scrounging (duration): feeding from a food box that another individual had opened
5. Observing others (duration): looking at an individual that successfully opened the food box by either technique or was manipulating the food box

### Behavioral observations

In addition, we conducted continuous focal animal observations that lasted 30 minutes per individual during the experimental period. Focal animals were selected in a random but balanced order. We recorded affiliative behaviors such as grooming, body contact and huddling according to an established ethogram (Pereira and Kappeler 1997).

### Statistical analysis

Model 1: To investigate factors predicting the probability of learning, we fitted a Bayesian logistic regression model with a Bernoulli error structure with a logit link function by using the brms package in R (Bürkner 2017). We included whether the individual learned to open the food box (yes or no) by either technique for each session until they opened the box for the first time as response. As fixed factors, we included the cumulative duration of manipulation of the food boxes and the cumulative duration of time spent watching successful individuals and scrounging across sessions until the first success as well as sex and age. Individual and group identity as well as the number of sessions were included as random factors. Time spent watching successful individuals correlated with time spent scrounging (Pearson correlation: R=0.74, *p*<0.001) but did not result in collinearity (vif=2.19). We specified weakly informative priors to regularize parameter estimates. Specifically, we assigned a normal (0, 2) prior to the regression coefficients (β) to allow for moderate variation in effect sizes while keeping the estimates centered around zero. For the standard deviations of the random effects, we used an exponential (1) prior, which constrains values to be positive while discouraging excessively large variance estimates.

Preferences of techniques: To determine whether the observed proportion of technique choice significantly differed from chance (prob=0.5), we used a one-sample proportion test. We calculated the proportion of using the lifting technique in relation to the total number of successful openings with both techniques to investigate whether the lifting technique was preferred over the pushing technique.

Model 2: To investigate if individuals became more efficient over time throughout the experimental sessions, we modeled the latency, i.e., time between the start of the manipulation to the successful opening of the food box, with a Linear Mixed Model (LMM). We included sex, age, used technique and total number of successes as fixed effects. Individual, group, and session identity were included as random intercept effects. To avoid overconfident estimates and keep the type I error rate at the nominal level of 0.05, we included random slopes of technique and number of successes within individual identity and of all fixed factors within group and session identity. We excluded the correlations among random slopes and intercepts due to convergence issues.

Model 3: To identify predictors of success, we fitted a Generalized Linear Mixed Model (GLMM) with a negative binomial error distribution. The response variable was the total number of successes per individual, with sex, age, and their interaction included as fixed effects, and group identity specified as random intercept effects. In addition, we included the log-transformed number of sessions individuals joined as an offset term. We included random slopes of fixed effects within group but we excluded the correlation among random slopes and intercepts due to convergence issues (Schielzeth and Forstmeimer 2009; Barr et al. 2013).

Model 4: In this GLMM with a Poisson distribution, we modeled the number of scroungers per event. We included the used technique, sex and age of the producer as fixed effects. The random intercept effects were producer and group identity; furthermore, we included the random slopes of technique and sex within producer as well as age within group. We removed the correlation between random intercepts and slopes due to convergence issues.

Model 5: We fitted a GLMM with a Poisson distribution to examine what predicts the frequency of scrounging for all individuals that participated in the experiment (N=24). We included the frequency of scrounging as response, whether they had learned to open the food box by themselves (as a binary variable: yes or no), sex, and age as fixed effects. Since individuals that spent more time at the food boxes had, in principle, more opportunities to scrounge than those that spent less time at the boxes, we included the time spent at the boxes as an additional control predictor. Group identity was included as random intercept effect. We included the random slopes of age and time spent at the boxes within group identity.

Model 6: Since individuals that have learned to open the boxes also scrounged (N=16), we examined whether success in opening the boxes predicts scrounging. We fitted a GLMM with a Poisson error structure modeling the frequency of scrounging. We included the number of successful openings and sex as fixed effects and group identity as random intercept effect, as well as the random slope of number of openings within group identity.

Models 7 and 8: We additionally investigated whether more successful individuals or individuals that were scrounged more often received more affiliation, i.e, time spent grooming, being in body contact and huddling. We calculated affiliation balance by subtracting the rate of affiliation given from the rate of affiliation received (in minutes) and dividing it by the total observation time (in hours) for all individuals. We used affiliation balance as the response variable in two linear mixed models (LMMs). In model 7, we included the log-transformed number of successes as a fixed effect. Additionally, we included the following control factors: sex, age, and participation (yes or no), to control for participation in the experiment; and a factor, "learning category". This factor controlled for the fact that individuals who learned the task earlier were able to perform more successful openings. The "learning category" factor comprised three levels: first and second learners (early learners), third and fourth learners (middle learners), and fifth and sixth learners (late learners). We included individual and group identity as random intercept effects, and the random slope of age and number of successes within group. We removed the correlation between random intercept and slope due to convergence issues. Model 8 was similar to model 7 but included the log-transformed frequencies of being scrounged as a fixed effect predictor instead of the number of successes and the random slope of age and frequencies of being scrounged within group.

### Model implementations

All analyses were conducted using R (v. 4.4.2; R Core team, 2024). To ease model convergence, we *z*-transformed continuous variables before including them in all models. To test the significance of fixed effect predictors as a whole, we compared the fit of the full model with that of the null model comprising only random factors (Schielzeth and Forstmeier 2009; Forstmeier and Schielzeth 2010). Additionally, we checked GLMMs for overdispersion by using the ‘DHARMa’ package (Hartig 2022). We obtained confidence intervals for all models by means of parametric bootstraps using the function ‘bootMer’ of the package ‘lme4’ (Bates et al. 2015), applying 1000 bootstraps. We checked for collinearity issues by determining variance inflation factors for a standard linear model without random effects using the package ‘car’ (Fox and Weisberg 2019). To estimate model stability, we dropped levels of the random effects one at a time from the dataset and compared the obtained estimates to the estimates obtained for the full dataset.

## Results

### Innovators and preferences for one of the two opening techniques

In total, 27 out of 29 individuals participated in the experiment, 24 interacted with the boxes, and 16 successfully opened a food box at least once using either technique. The first individuals learning how to open the food boxes successfully with either technique in each group were two juvenile females, one juvenile male and an adult female. In two groups, innovators for each technique were different individuals (one adult female, one juvenile female, and two juvenile males) whereas for the other two groups, both techniques were discovered for the first time by the same individuals (one adult female and one juvenile female). The first success of innovators for each technique varied between 1^st^ session to 4^th^ session out of 10.

Out of 16 individuals, eight individuals used the pushing and eight individuals used the lifting technique for the first successful opening. Eight individuals did not show flexibility and used the same technique throughout the experiment (three pushing and five lifting), whereas the other eight individuals used both techniques. The result of a one-sided proportion test indicates a strong preference for the lifting technique, with a significantly greater proportion of individuals - ten using mostly the lifting technique, four using mostly the pushing technique, and two using both techniques equally across sessions - preferring the lifting technique over an equal preference baseline (one sample proportion test, prob,>0.5; observed prob.=0.8165; χ^2^(1)=522.88, *p*<0.001, 95% CI [0.798, 1.000]).

### What predicts initial learning: use of individual or social information?

The Bayesian logistic regression (Model 1) revealed that the probability of learning was predicted by time spent manipulating the food box (Table 1), but not by the time spent watching successful individuals or scrounging. Similarly, age and sex did not predict learning (Table 1). Thus, individuals who manipulated the food boxes for longer were more likely to learn the task. The model showed good convergence (Rhat = 1.00 for all parameters), and effective sample sizes were sufficient (Bulk_ESS > 1500, Tail_ESS > 1500).

**Table 1:** Results of the Bayesian logistic regression on the effects of scrounging and individual information on the probability of learning the task.

| Model | Predictor | Estimate | SE | Lower<br>95% CI | Upper<br>95% CI |
| --- | --- | --- | --- | --- | --- |
| Model 1:<br>Probability of<br>learning | Intercept | -2.49 | 1.18 | -4.98 | -0.18 |
|  | Time spent manipulating | 2.63 | 0.84 | 1.24 | 4.55 |
|  | Time spent scrounging | 0.75 | 0.87 | -0.92 | 2.53 |
|  | Time spent watching<br>successful individuals | -0.79 | 1.11 | -3.00 | 1.44 |
|  | Sex (male) | 0.33 | 1.23 | -2.12 | 2.64 |
|  | Age | -0.34 | 0.68 | -1.85 | 0.91 |

**Table 2:** Results of a) the LMM on the effects of sex, age, used technique and success on the latency to succeed and b) the GLMM on the influence of sex and age on the success.

| Model | Predictor | Estimate | SE | *p*-value |
| --- | --- | --- | --- | --- |
| a) Model 2: Latency to open the food box | Intercept | 2.667 | 5.254 | <sup>1</sup> |
|  | Sex (male) | 6.349 | 6.843 | 0.393 |
|  | Age | -2.855 | 6.268 | 0.656 |
|  | Technique (slide) | 14.345 | 17.486 | 0.429 |
|  | Total success | 0.797 | 0.563 | 0.267 |
| b) Model 3: Success | Intercept | 2.105 | 0.298 | <sup>1</sup> |
|  | Sex (male) | -1.946 | 0.543 | 0.001 |
|  | Age | 0.011 | 0.250 | 0.965 |
|  | Sex*age | -3.198 | 0.691 | 0.001 |

### Who scrounges from whom?

Individuals scrounged on average in 26.12 ± 22.26% of successful openings. On average 1.30 ± 0.62 (mean ± SD, range: 1-5) individuals scrounged per event. The number of scroungers did not differ depending on the used techniques (lifting or pushing), sex and age of the producer (Table 3a, likelihood ratio test: full – null model comparison: χ^2^=1.30, df=3, *p*=0.729).

**Table 3:** Results of the GLMMs estimating a) the effect of technique, sex and age of the provider on the number of scroungers per event, b) the influence of sex, age, and time spent at the feeding boxes on the frequency of scrounging, c) the effect of number of successful openings and age on the frequency of scroungers for learners only, the influence of d) success or e) time being scrounged and learning order (early, middle, late learners) on affiliation balance.

| Model | Predictor | Estimate | SE | P-value |
| --- | --- | --- | --- | --- |
| a) Model 4: Number of scroungers per event | Intercept | 0.309 | 0.118 | <sup>1</sup> |
|  | Technique (push) | -0.705 | 0.129 | 0.583 |
|  | Sex provider (male) | -0.013 | 0.114 | 0.908 |
|  | Age provider | -0.017 | 0.020 | 0.400 |
| b) Model 5: Frequency of scrounging including learner and non-learner | Intercept | 0.912 | 0.460 | <sup>1</sup> |
|  | Learned (yes) | 0.931 | 0.201 | 0.001 |
|  | Sex (male) | 1.030 | 0.136 | 0.001 |
|  | Age | -0.832 | 0.905 | 0.381 |
|  | Time at box | 0.203 | 0.110 | 0.064 |
| c) Model 6: Frequency of scrounging including only learner | Intercept | 1.222 | 1.468 | <sup>1</sup> |
|  | Success | -3.207 | 1.289 | 0.013 |
|  | Sex (male) | -0.007 | 0.158 | 0.964 |
|  | Success*sex (male) | 0.599 | 0.204 | 0.003 |
| d) Model 7: Influence of success on affiliation balance | Intercept | -10.903 | 2.622 | <sup>1</sup> |
|  | Success | 4.725 | 1.718 | 0.022 |
|  | Sex (male) | 1.401 | 1.898 | 0.464 |
|  | Age | 0.287 | 0.192 | 0.144 |
|  | Learning category (middle) | 2.801 | 2.476 | 0.264 |
|  | Learning category (late) | 8.111 | 3.452 | 0.024 |
|  | Participate (yes) | -0.748 | 2.637 | 0.778 |
| e) Model 8: Influence of time being scrounged on affiliation balance | Intercept | -11.533 | 3.102 |  |
|  | Frequency being scrounged | 4.789 | 2.015 | 0.044 |
|  | Sex (male) | 1.294 | 2.011 | 0.524 |
|  | Age | 0.360 | 0.205 | 0.087 |
|  | Learning category (middle) | 4.419 | 2.859 | 0.130 |
|  | Learning category (late) | 8.284 | 3.834 | 0.037 |
|  | Participate (yes) | -0.511 | 2.727 | 0.852 |

Three individuals that participated in the experiment never scrounged or opened the food boxes, and only five individuals relied solely on scrounging. The frequency of scrounging was higher in learners than in non-learners and higher in males compared to females (likelihood ratio test: full – null model comparison: χ^2^=94.096, df =3, *p*<0.001, Table 3b, Figure 3a, b). However, age or the time spent exploring and manipulating (Table 3b).

**Figure 3:**
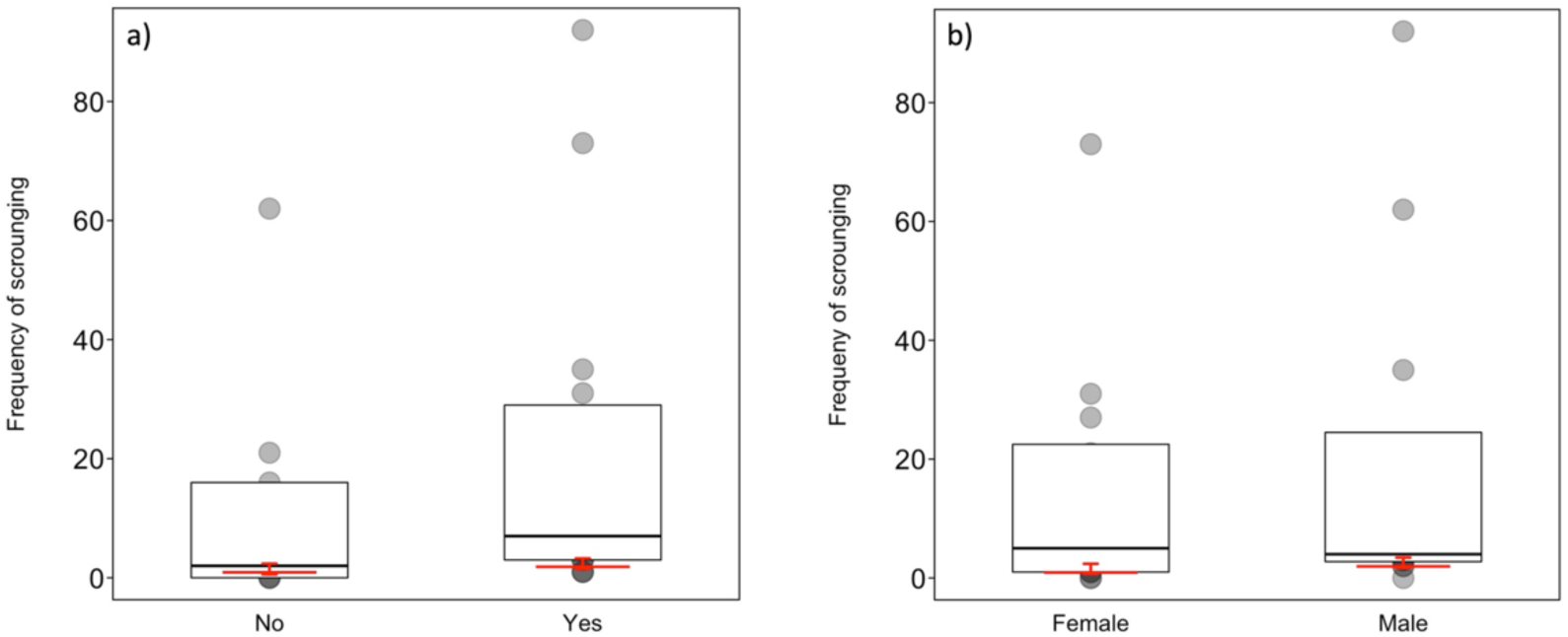
Frequency of scrounging as a function of a) whether individuals learned to open the feeding boxes or not and b) sex. Indicated are the median (horizontal lines) and quartiles (boxes). The red crosses represent the fitted model and its confidence limits (conditional on all covariates and factors centred to a mean of zero).

**Figure 4:**
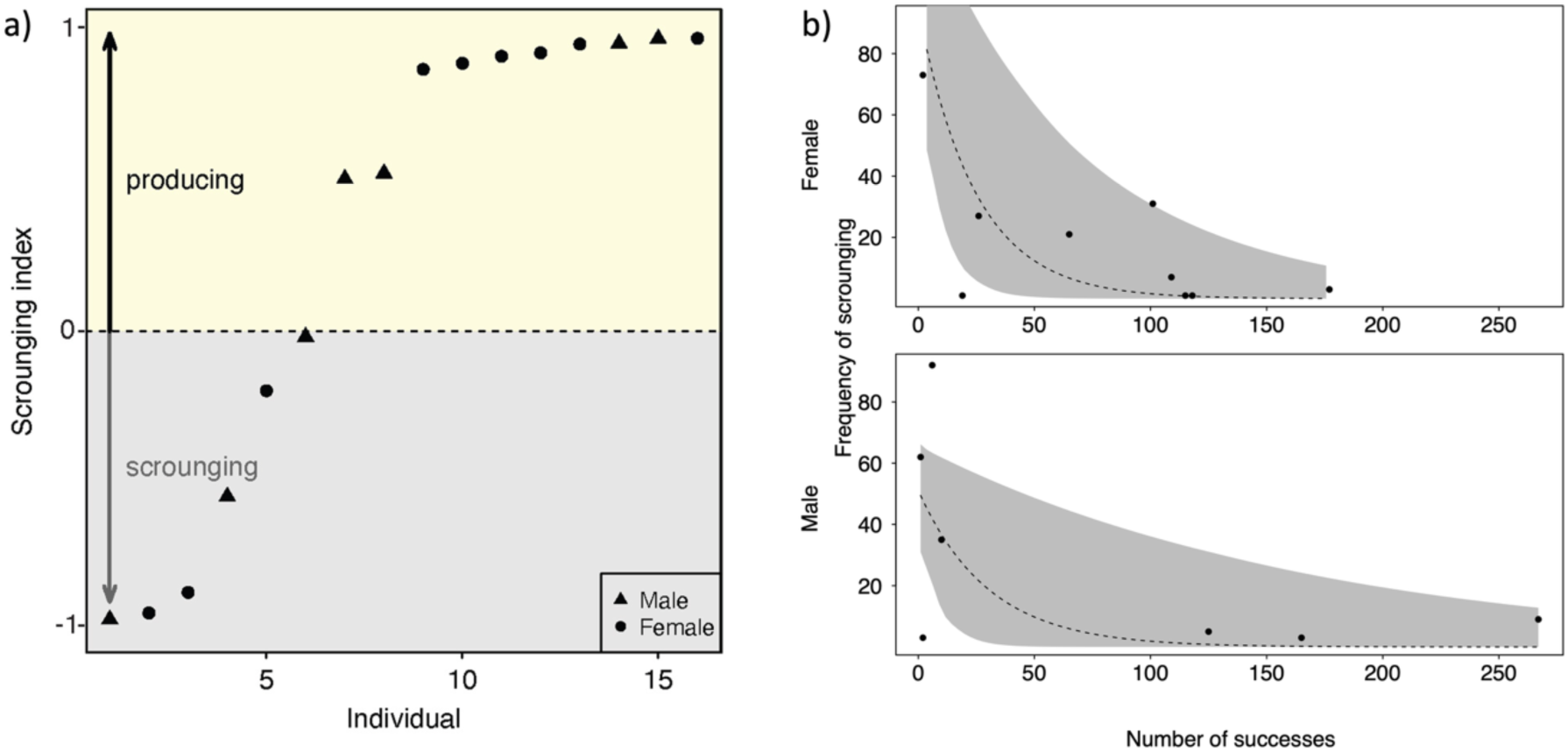
a)Scrounging index (N successful opening – N scrounging events divided by the sum of both) for each learner (N=16) and b) frequency of scrounging as a function the number of successful openings for learners only. The dashed line represents the regression lines and the shaded areas the 95% intervals.

All individuals that learned to open the boxes also scrounged, suggesting that they actively switched tactics. There was a strong variation in using either tactic, with some individuals relying mainly on scrounging and others mainly relying on producing (Figure 5a). We found that the interaction between success and age was significant, with less successful producers scrounging more often, and this effect was more pronounced in males (likelihood ratio test: full – null model comparison: χ^2^=16.867, df=3, *p*<0.001, Table 3c, Figure 5b).

**Figure 5:**
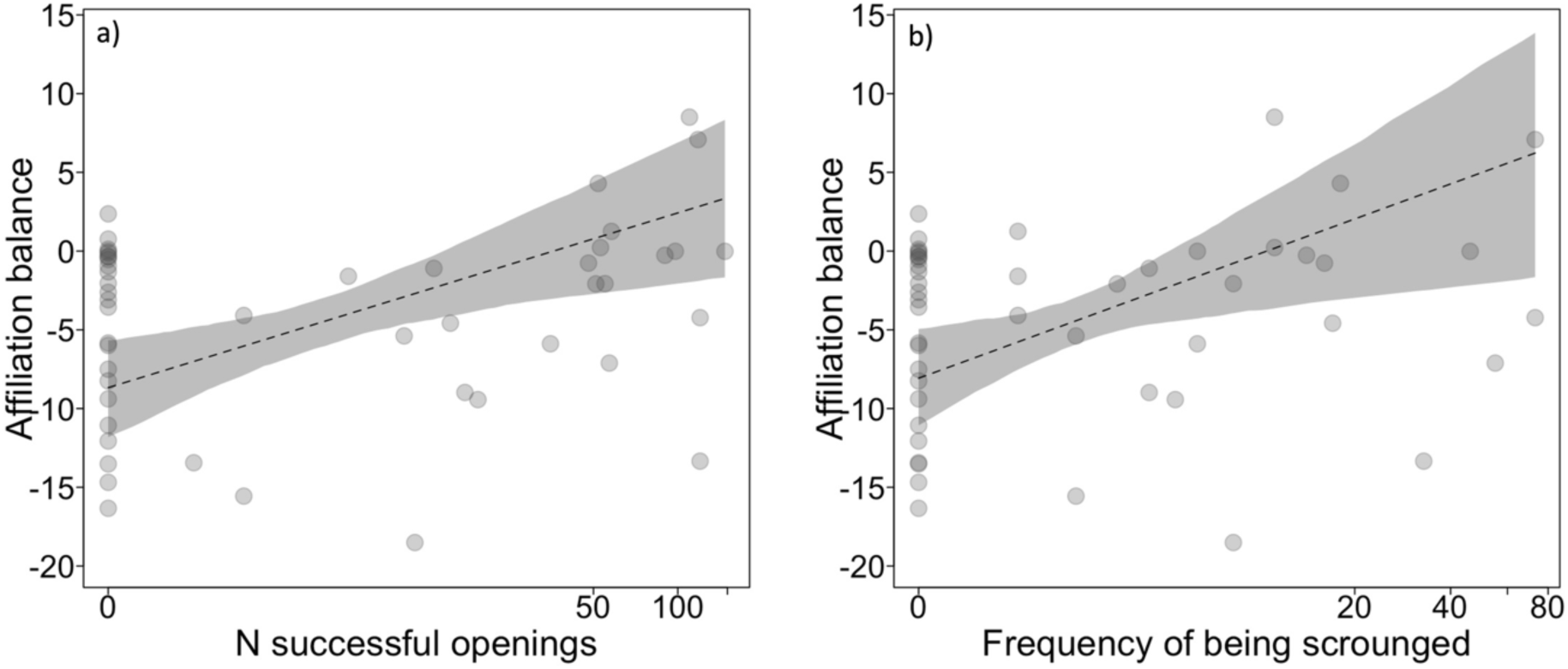
Affiliation balance as a function of the number of a) successful openings and b) the frequency of scrounging. The dotted lines indicate the regression lines and grey polygons present the 95^th^ intervals.

### Links between affiliative relationships, success, and scrounging

We found that more successful individuals received more affiliation from group members than they gave and also late learners compared to first learners (Figure 5a, Table 3d, likelihood ratio test: full – null model comparison: χ^2^=6.789, df=1, *p*=0.009). Sex, age, and participation (yes or no) in the experiment did not influence affiliation balance (Table 3d). Since the frequency of being scrounged correlated positively with the number of successful openings (Pearson’s correlation: R=0.69, *p*<0.001, N =16), we estimated in an additional model whether individuals that were scrounged more often also received more affiliation. We found that individuals that were victims of scrounging received more affiliation, but this effect was weaker than for the success model (Figure 5b, Table 3e, likelihood ratio test: full – null model comparison: χ^2^=5.931, df=1, *p*=0.015). Also, late learners received more affiliation compared to middle learners but sex, age, and whether they participated in the experiment or not did not influence affiliation balance (Table 3e).

## Discussion

In this study, we examined the interaction between social learning and scrounging in an open diffusion food box experiment and its relationship with social relationships in wild redfronted lemurs. Three out of the four innovators were juveniles, though age did not directly affect learning likelihood. Time spent manipulating boxes, and not time spent watching successful individuals or scrounging, predicted learning. Opening efficiency did not improve over time. With increasing age males were less successful compared to females. Scrounging frequency per event was not influenced by producer’s age, sex or technique used. Independent of age, learner and males scrounged more often than non-learners and females. Among learners, less successful individuals scrounged more often, and this effect was more pronounced in males. Most interestingly, more successful individuals received more affiliative behavior as well as individuals that were scrounged more often, though the latter effect was weaker. Thus, cognitive competence and scrouging tolerance may enhance social relationships in this tolerant primate species. We discuss these findings in detail below.

### Innovators and preferences for either technique

Innovators were mainly juveniles and females, aligning with previous findings in the same population of redfronted lemurs (Schnoell and Fichtel 2012). Since three out of four innovators were juveniles, this may suggest that juveniles have more free time to explore and thus greater opportunities to innovate (Kummer and Goodall 1985). Alternatively, their lower competitive abilities compared to adults might force them to access resources in novel ways (Laland and Reader 1999). Similarly, several other studies reported that young primates are more innovative than their older conspecifics, most likely because juveniles exhibit stronger exploratory and risk-taking tendencies than adults (Boesch and Boesch 1981; Cambefort 1981; Fragaszy and Adams-Curtis 1991; Hannah and McGrew 1987; Hauser 1988; Huffman 1996; Itani 1965; Kawamura 1954; Kummer 1971; Kummer and Goodall 1985; McGrew et al. 1979; Nishida et al. 1983). In contrast, a review study suggested the opposite trend, that innovators are typically adult males (Reader and Laland 2001). A possible explanation is that males, typically higher in the dominance hierarchy due to sexual size dimorphism, often monopolize food sources, providing them with more opportunities to innovate. Furthermore, larger body size in some species may lead to greater energetic needs (Bean 1999), potentially motivating males to explore novel foraging strategies more than females (Reader and Laland 2001). However, since redfronted lemurs exhibit no sexual dimorphism (Kappeler 1991) and live in a matrilineal and egalitarian society (Ostner and Kappeler 1999; 2004), our findings point to two main factors driving innovations in this species: (1) the availability of spare time and the risk-taking tendencies of juveniles, and (2) the lack of sexual size dimorphism, which likely enables females to access foraging opportunities more readily. The latter aligns with observations in other lemur species (Kappeler 1987; Schnoell and Fichtel 2012; Huebner and Fichtel 2015, but see Dean et al. 2011).

In addition, we found that individuals exhibited flexibility in their choice of technique during the initial trial, contrasting with the findings from a previous social learning experiment involving trained demonstrators (Schnoell and Fichtel 2012). Similarly, wild jackdaws (*Corvus monedula*) exhibited flexibility in food preferences of novel food in the absence of a demonstrator in an open-diffusion paradigm (Arbon et al. 2017). In our study, some individuals developed a consistent preference for one technique over the other during the experimental sessions, with ten favoring the lifting technique and four favoring the pushing technique. However, since technique choice did not influence the likelihood of being scrounged from, these preferences cannot be attributed to attempts to avoid scroungers. Instead, the observed bias in technique use may reflect the tendency for arbitrary traditions to emerge more easily in small groups (Franz and Matthews 2010). It is possible that an initial slight preference for the lifting technique was amplified through social facilitation and individual reinforcement, leading it to become both the most commonly used and the preferred technique among most individuals. However, additional network biased diffusion analyses are indicated to confirm this assumption (Canteloupe et al. 2020; Hasenjager et al. 2021).

### What predicts initial learning: use of individual or social information?

The probability of learning to open the food box was predicted by use of individual information, i.e., the time spent manipulating the boxes, but not by use of social information, i.e., the time spent watching successful individuals or scrounging. Hence, scrounging did neither hinder nor facilitate initial learning in redfronted lemurs. In principle, scrounging can be used to gather social information on how to solve the task as suggested for common marmosets (*Callithrix jacchus*, Caldwell and Whiten 2003). Scrounging can also inhibit social learning or learning in general (Giraldeau and Lefebvre 1987; Beauchamp and Kacelnik 1991; Reichert et al. 2021; Lefebvre and Helder 1997). For example, scroungers tended to learn a task more slowly in great tits (*Parus major;* Aplin and Morand-Ferron 2017) or in mixed-species flocks of great and blue tits (*Cyanistes caeruleus;* Reichert et al. 2021). However, as we only investigated the influence of scrounging on the first, initial learning success, we cannot rule out the possibility that social information gathered during scrounging or the time spent watching successful individuals is related to the preferences of the technique used within groups. As mentioned above, since some individuals preferentially used one of the two techniques, further analyses are required to investigate whether the spread of using a specific technique is associated with the scrounging network. In addition, the time spent watching successful individuals did not predict initial success. Since the time spent watching successful individuals correlated with the time spent scrounging, it is possible that scroungers watched successful individuals to seek scrounging opportunities, which is difficult to demonstrate.

### Efficiency and success in opening the food boxes

The latency to open a food box did not decrease over the course of the experiment, regardless of the technique used, suggesting that no additional learning was required to open the boxes more efficiently and that both techniques may have been equally difficult to open. Females were more successful than older males, consistent with earlier studies on redfronted lemurs (Schnoell and Fichtel 2012) and ring-tailed lemurs (Kappeler 1987). In another open diffusion task in wild vervet monkeys, males and higher-ranking individuals were more successful than lower ranking individuals and females (Canteloup et al. 2021). Since vervet monkeys exhibit sexual size dimorphism, with males being bigger than females and a propensity to dominate females (Saccà et al. 2022), the variation in success might be due to a greater potential to monopolize access to food (Canteloupe et al. 2021). However, lemurs do not exhibit sexual size dimorphism, while females are dominant over males in ring-tailed lemurs but not in redfronted lemurs (Kappeler et al. 2022). Thus, the female-biased success in ring-tailed lemurs might be due to the monopolization potential of dominant females. However, the age-related higher success of females in redfronted lemurs cannot be explained by a dominance-mediated higher monopolization potential.

### Who scrounges from whom?

Scrounging occurred in about a quarter of successful openings with one to five individuals scrounging from the same box at a time. Scrounging was not influenced by producer’s age, sex or technique used. Only five individuals relied solely on scrounging, while learners scrounged more often than non-learners. Thus, scrounging cannot be considered as an alternative strategy to producing. Males who were overall less successful at opening the boxes than females also scrounged more often. Individuals that learned to open the food boxes by themselves actively switched between scrounging and producing. Less successful individuals scrounged more often and this effect was more pronounced in males. In some species it has been suggested that dominant or more competitive individuals scrounge more often because they are better at displacing others at food resources (Lendvai et al. 2006; Lee and Cowlishaw 2017; Cram et al. 2023). In contrast, in rooks (*Corvus frugilegus*), dominant individuals produced more often because they were able to monopolize the food resource, but mixed tactics of producing and scrounging were guided by tolerance among paired birds (Jolles et al. 2013). In our experiment, on average more than one redfronted lemurs scrounged simultaneously. As redfronted lemurs do not exhibit clear dominance relationships (Ostner and Kappeler 1999), but rather high levels of social tolerance, with individuals feeding in close proximity at a valuable food resource with little or no aggression (Fichtel et al. 2018), this social tolerance may explain the observed scrounging pattern.

### Links between affiliative relationships, success, and scrounging

In this study, we found that more successful redfronted lemurs received more affiliation from other group members. These findings are in line with other studies showing that skilled individuals become more valuable social partners when they possess new information (Kulahci et al. 2018) or provide food that allows others to scrounge (Stammbach 1988; Fruteau et al. 2009; O’Hearn et al. 2025). In our study, we also considered whether affiliation is traded as a commodity to scrounge. More successful individuals were scrounged from more often. Producers also received more affiliation, but the effect was stronger for successful individuals. Hence, further research is needed to disentangle whether individuals traded affiliation for proximity to individuals to gather social information or for tolerance to scrounge. Finally, last learners were given more affiliation in comparison to first learners, which is difficult to explain

## Conclusions

Understanding how individuals navigate complex social interactions and exploit social information provides insights into the evolutionary significance of behavioral flexibility and social competence (Kulahci and Quinn 2019; Cram et al. 2023). Our findings highlight the complex interplay between social behavior and cognitive performance in redfronted lemurs, where manipulation success—not observation or scrounging—best predicted learning. Social rewards were directed toward both successful individuals and those who tolerated scrounging, suggesting that cognitive competence and social tolerance for scroungers may enhance social relationships in this tolerant primate species.

## Acknowledgments

We are grateful to our team in Madagascar for support during this study, and Roger Mundry for statistical advice. We also thank the MEDD Madagascar, the MZBA of the University of Antananarivo, and the CNFEREF Morondava for authorizations of this study. This publication was funded by the Deutsche Forschungsgemeinschaft (DFG, German Research Foundation) - Project-ID 454648639 - SFB 1528 – Project B06.

## Author contributions

CF, PMK, EK conceived the study; RV, MM, JP-C conducted the study; EK, CF, RV, MM, JP-C analyzed the data; EK, CF drafted the MS, and all authors reviewed the MS.

## Data availability

The data of this study have been deposited in https://figshare.com/s/73e910a99db9a6005693

## Declarations

### Ethical approval

This study adhered to the ASAB/ABS Guidelines for the Treatment of Animals in Behavioural Research and Teaching and to the legal requirements of the country (Madagascar) in which the study was carried out. The protocol for this research was approved by the Malagasy Ministry of the Environment, Water, and Forests (064,22/ MEDD/ SG/ DGGE/ DAPRNE/ SCBE.Re).

### Consent to participate

N/A.

